# A deep generative model for estimating single-cell RNA splicing and degradation rates

**DOI:** 10.1101/2023.11.25.568659

**Authors:** Chikara Mizukoshi, Yasuhiro Kojima, Satoshi Nomura, Shuto Hayashi, Ko Abe, Teppei Shimamura

**Affiliations:** Division of Systems Biology, Graduate School of Medicine, Nagoya University, Aichi, Japan; Nagoya University Hospital, Aichi, Japan; Laboratory of Computational Life Science, National Cancer Center Research Institute, Tokyo, Japan; Division of Computational and Systems Biology, Medical Research Institute, Tokyo Medical and Dental University, Tokyo, Japan

**Keywords:** single-cell RNA sequencing (scRNA-seq), RNA splicing, RNA degradation, splicing kinetics, transcriptome dynamics, RNA-binding proteins, RNA velocity, neural network, variational autoencoder (VAE), deep generative model, dimensionality reduction, cell differentiation, metabolic labeling

## Abstract

Messenger RNA splicing and degradation are critical for gene expression regulation, the abnormality of which leads to diseases. Previous methods for estimating kinetic rates have limitations, assuming uniform rates across cells. We introduce DeepKINET, a deep generative model that estimates splicing and degradation rates at single-cell resolution from scRNA-seq data. DeepKINET outperformed existing methods on simulated and metabolic labeling datasets. Applied to forebrain and breast cancer data, it identified RNA-binding proteins responsible for kinetic rate diversity. DeepKINET also analyzed the effects of splicing factor mutations on target genes in erythroid lineage cells. DeepKINET effectively reveals cellular heterogeneity in post-transcriptional regulation.

## Introduction

Messenger RNA (mRNA) splicing and degradation play essential roles in precise gene expression regulation. These processes are vital for accurate utilization of genetic information within cells. Inappropriate splicing can lead to production of dysfunctional proteins, potentially resulting in severe implications for fundamental cellular functions. Recent studies have established that abnormal mRNA splicing and degradation are closely associated with development and progression of diseases such as cancer [Bradley *et al*., 2023, Fang et al., 2022].

Several methodologies are available to estimate mRNA splicing and degradation rates, each with its own limitations and challenges. Metabolic labeling methods [Battich *et al*., 2020, Qiu *et al*., 2020] are used to estimate the synthesis and degradation rates in genome-wide RNA metabolism by integrating RNA metabolic labeling with cell-specific splicing kinetics. However, owing to the necessity of specific metabolic labeling, this approach is limited and cannot be applied as readily as conventional scRNA-seq data. Combination of scRNA-seq data with the RNA velocity theory [La Manno *et al*., 2018] was introduced to model the dynamic processes of mRNA in individual cells. However, this approach has been criticized for assuming uniform kinetic rates across cells, which may cause misrepresentation of true biological variation. While transcription rates have been modeled to account for cell-to-cell variability, methods such as scVelo [Bergen *et al*., 2020] and VeloVI [Gayoso *et al*., 2024] have assumed uniform splicing and degradation rates for each gene. A novel relay velocity model [Li *et al*., 2023] utilizes neighboring cell information and leverages deep neural networks to estimate cell-specific kinetic rates. However, its primary intention is to refine the RNA velocity, leaving questions regarding the accuracy of the kinetic rates for each cell.

In light of these challenges, we introduced DeepKINET (a deep generative model with single-cell RNA kinetics), an advanced analysis framework based on deep generative modeling. This framework uses deep generative model-driven cell states in scRNA-seq data to accurately estimate single-cell splicing and degradation kinetics. Our method aims to reveal the heterogeneity in splicing and degradation rates across cells, enabling to elucidate post-transcriptional regulatory mechanisms mediated by factors such as RNA-binding proteins.

We demonstrate that DeepKINET can estimate mRNA splicing and degradation rates with greater precision than existing methods, as evidenced by simulated and metabolic labeling experimental data. Moreover, we demonstrate its robustness against dropouts. By applying DeepKINET to a forebrain dataset, we analyzed whether genes governed by the same RNA-binding proteins have equivalent trends in their splicing and degradation rates, and we identified the biological functions of these RNA-binding proteins. Furthermore, when applied to breast cancer data, DeepKINET revealed splicing and degradation anomalies related to cancer metastasis; we provide specific examples. In addition, we analyzed the effects of mutations in a splicing factor on the target genes in erythroid lineage cells. The results enhance our understanding of mRNA splicing and degradation processes and help to elucidate underlying molecular mechanisms and potential therapeutic targets.

## Results

### Conceptual view of DeepKINET

Figure 1 presents a clear overview of the conceptual framework of DeepKINET. This method processes both spliced and unspliced mRNA counts from scRNA-seq data and subsequently generates comprehensive kinetic rates across genes, including splicing and degradation rates, at single-cell resolution. DeepKINET addresses heterogeneity in kinetic rates spanning genes and cells, which is ignored by existing methods [La Manno *et al*., 2018, Bergen et al., 2020].

**Figure 1:**
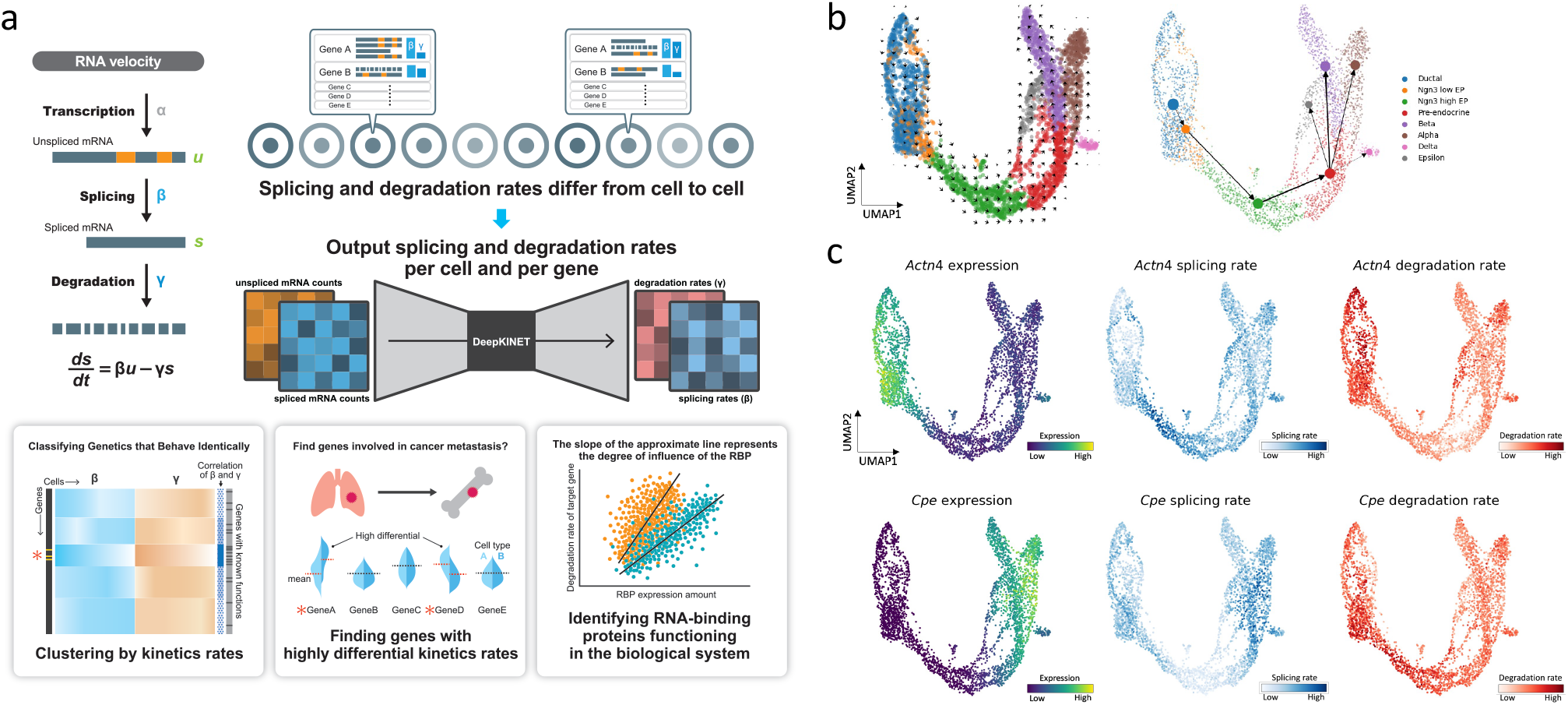
Overview of DeepKINET. **a**. Overview of our method for estimating single-cell transcriptome dynamics from latent variables. DeepKINET receives scRNA-seq data that have unspliced and spliced counts and outputs kinetic rates at the single-cell level. DeepKINET provides biologically meaningful insights by accounting for cellular heterogeneity in kinetic rates, which is ignored by existing methods. For example, DeepKINET can be used to classify genes by their kinetic rates, find genes that show significant rate variation among cell populations, and identify RNA-binding proteins involved in splicing and degradation. **b**. Estimated RNA velocity by DeepKINET in the mouse pancreas dataset visualized on Uniform Manifold Approximation and Projection (UMAP) embedding. The direction of transition in latent space is plotted in 2D coordinates in the same way as scvelo. Trajectory inference by PAGA[Wolf *et al*., 2019] was performed using RNA velocity from DeepKINET. **c**. Expression, splicing rate and degradation rate at the single-cell level projected on UMAP embedding. DeepKINET estimates splicing and degradation rates for each cell based on the RNA velocity equation and cell states. The colors of the points indicate the gene expression, the splicing rate, and the degradation rate per cell.

DeepKINET uses a deep generative model of mature and immature transcripts based on an RNA velocity equation. This enables optimization in which the splicing and degradation rates are adjusted according to the cell state. First, we use a variational autoencoder (VAE) to model stochastic transitions within the latent cell state space, similar to that in our previous study [Nagaharu *et al*., 2022]. DeepKINET assumes that the kinetic parameters for each cell are obtained from transformation of the latent cell state by the neural network. We optimized both cell state dynamics and kinetic parameters to align with the observed mature and immature transcript levels, following the RNA velocity equation.

Beyond kinetic rate heterogeneity estimation across genes and cells, DeepKINET offers the following: 1. Gene clustering based on kinetic rates, which enables identification of genes with analogous rate patterns; 2. Identification of genes exhibiting significant rate variations by comparing different cell populations; and 3. Detection of RNA-binding proteins that influence splicing and degradation rates of their associated targets.

DeepKINET not only delivers refined insights into RNA kinetics, but also serves as a springboard for in-depth molecular studies, promising deeper comprehension and demystification of the complex regulatory mechanisms guiding cellular kinetics. It is accessible as a user-friendly open-source Python package with comprehensive documentation at https://github.com/3254c/DeepKINET.

### Simulated data to demonstrate accuracy and superiority of DeepKINET

We used simulated data to evaluate the accuracy of the kinetic rates estimated using the DeepKINET software. Simulated data were generated using SERGIO [Dibaeinia *et al*., 2020], which uses gene regulatory networks and RNA velocity equations to generate the scRNA-seq data. We generated scRNA-seq count data for each cell cluster with different splicing and degradation rates.

We applied DeepKINET to each simulated dataset and confirmed that it predicted the correct direction of differentiation (Fig. S1a). We then estimated the kinetic rates for each single cell (Fig. 2a), averaged them over each cell cluster, and calculated the correlation coefficient using the set value (Fig. 2b). We found positive correlations across various dropout scenarios. Therefore, we concluded that our method is robust against data sparsity. The existing method, cellDancer [Li *et al*., 2023], showed positive correlations in splicing rates, whereas DeepVelo [Cui *et al*., 2024] showed negative correlations in splicing rates. Both methods were less accurate than DeepKINET. Furthermore, cellDancer showed negative correlations in degradation rates. DeepVelo showed positive correlations with respect to degradation rates, but DeepKINET had higher correlations. Subsequent simulations were conducted using varying numbers of cells. For these simulations, we used the default dropout rates. We applied DeepKINET to each simulated dataset and computed correlation coefficients for the set values. DeepKINET could accurately estimate the kinetic rates, even for small numbers of cells (Fig. 2c). On the other hand, cellDancer also showed positive correlations in splicing rate estimation accuracy, but it was less accurate than DeepKINET and required more cells until the estimation accuracy stabilized. Furthermore, DeepVelo was unable to make accurate estimates even when the number of cells was increased. In degradation rates, cellDancer consistently failed to make correct estimates, and the accuracy decreased as the number of cells increased. DeepVelo showed positive correlations, but the estimation precision was still lower than DeepKINET.

**Figure 2:**
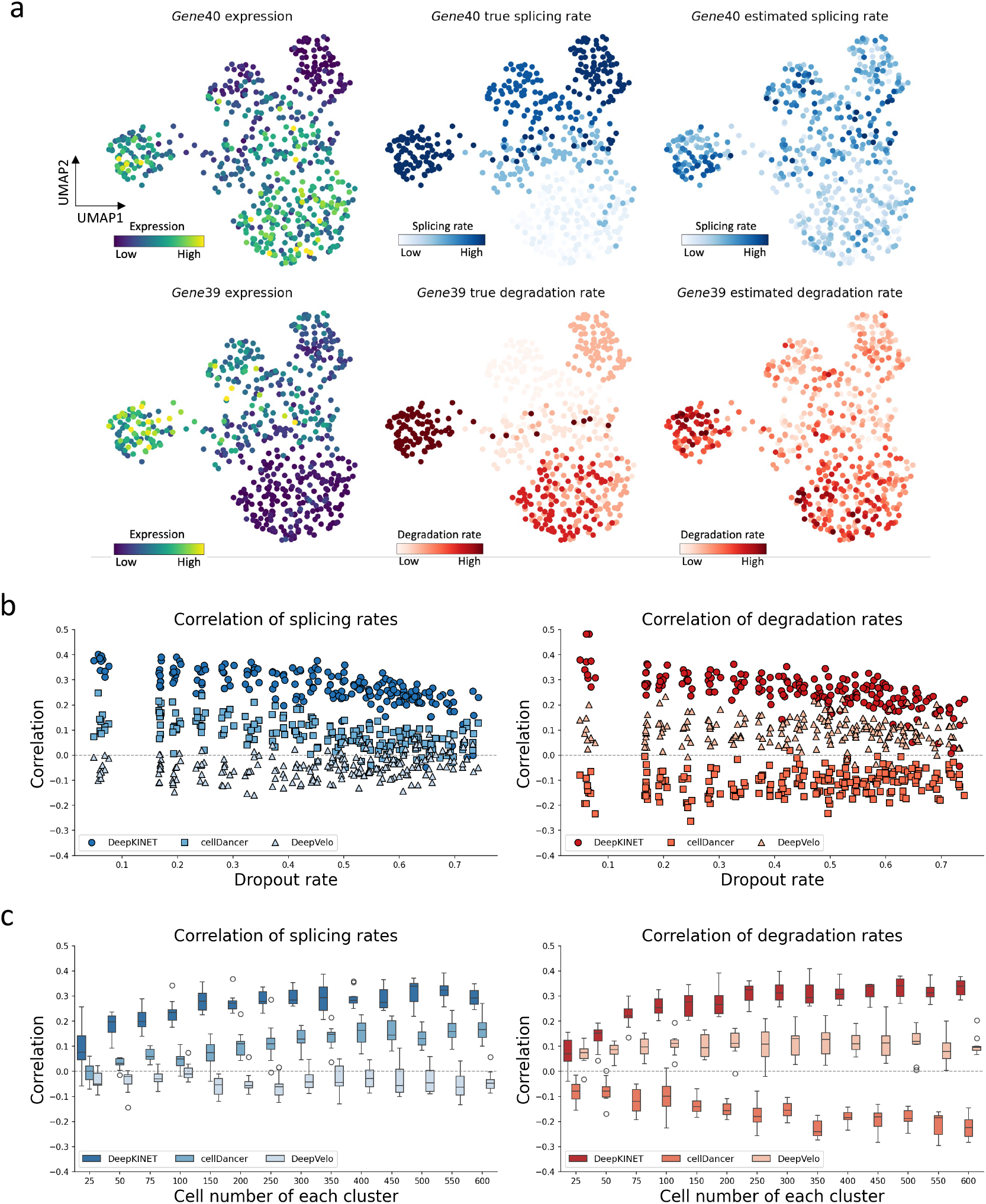
DeepKINET is robust to dropout rates and cell numbers in simulated data, and its performance exceeds that of cellDancer and DeepVelo. **a**. Visualization of the UMAP embedding of the expression, set kinetic rates, and estimated kinetic rates. The gene with the highest correlation in splicing rate and the gene with the highest correlation in degradation rate are shown. To prevent extreme values from affecting the visualization, the minimum or maximum value of the top 1% was forced to the 1% and 99% quantile values. **b**. Scatter plot of correlation coefficient averages of splicing rates and degradation rates for each dataset. Ten datasets were generated for each of the 20 different generation conditions. We applied DeepKINET, cellDancer and DeepVelo once to each dataset and calculated the correlation coefficient between the set rates and the estimated rates by each method. DeepKINET’s accuracy exceeds that of cellDancer and DeepVelo. **c**. Box plot of correlation coefficient averages when varying the number of cells in a cluster. Ten datasets were generated for each of the 14 different generation conditions. DeepKINET always had a positive correlation coefficient and outperformed cellDancer and DeepVelo.

These validations confirmed the accuracy of the splicing and degradation rates estimated by DeepKINET, marking a clear advancement over the kinetic parameter estimation capabilities of cellDancer and DeepVelo. Notably, the accuracy of splicing rate estimation by cellDancer appeared to increase slowly as the number of cells increased, implying a requirement for larger datasets than those required by DeepKINET for accurate predictions. A detailed exposition of genes that were successfully estimated and those that were not is shown in Supplementary Figure 1.

Due to the presence of multiple unknowns in the RNA velocity equation for spliced mRNA, the solution of the splicing and degradation rates for each cell may be underdetermined, which could lead to correlation between the estimated kinetic rates. To assess whether each method can estimate these parameters independently, we investigated the correlation between the estimated splicing rates and degradation rates in each simulation dataset. We observed that the correlation between splicing and degradation rates estimated by DeepKINET was lower than that of cellDancer and DeepVelo (Fig. S1d). This suggests that DeepKINET can estimate splicing and degradation rates more independently compared to other methods. In contrast, cellDancer and DeepVelo exhibited relatively high correlations, indicating that these methods have difficulty in separately considering splicing and degradation processes.

### Accuracy of DeepKINET for real data evaluated using metabolic labeling data

We next evaluated the accuracy of DeepKINET for real data using multicellular-level kinetic rates derived from metabolic labeling experimental data. The values obtained from the metabolic labeling experiments depended on the assumptions of the mathematical model used and did not represent the perfect ground truth. Nevertheless, the temporal resolution inherent in the metabolic experimental data lost in scRNA-seq provides a benchmark from which to assess the similarity to extrapolated kinetic rates. Li et al.[Li *et al*., 2023] used single-cell EU-labeled RNA sequencing (scEU-seq) [Battich *et al*., 2020] to qualitatively assess the accuracy of cellDancer, which is limited to cell cycle genes.

Using the same scEU-seq cell cycle dataset, we evaluated the accuracy of DeepKINET. scEU-seq methodology can be used to estimate multicellular-level kinetic rates by observing temporal variations in the fraction of 5-ethynyl-uridine(EU)-labeled mRNA. Battich et al. did not differentiate between unspliced and spliced mRNAs when modeling mRNA metabolism. Conversely, Dynamo [Qiu *et al*., 2022] can estimate kinetic rates, including splicing rates, by accounting for splicing events in the scEU-seq data. We partitioned the cell cycle dataset into PULSE and CHASE experimental categories, each distinctly modeling mRNA metabolism. In the Pulse experiment, the EU incubation time differs for each cell. In the Chase experiment, EU incubation is performed under the same conditions, followed by washing time with uridine, which varies depending on the cells. We divided the cells into six to thirty clusters, each containing an equal number of cells across the cell cycle trajectory. Dynamo was used to determine the splicing and degradation rates for each cluster.

Next, we estimated the RNA velocity using DeepKINET and confirmed that the estimated future states of individual cells followed the order of the cell cycle (Fig. 3a, S2a). We then estimated the single-cell splicing and degradation rates, averaged them across clusters, and calculated the correlation coefficient using the kinetic rates determined using Dynamo. Our method showed positive correlations in both PULSE and CHASE experiments, outperformed cellDancer and DeepVelo in terms of accuracy in degradation rates, and demonstrated comparable performance in accuracy in splicing rates (Figure 3b, S2b). Notably, the PULSE experimental data were considered more reliable because the proportion of cells in different cell cycles was constant. Regarding the degradation rates, cellDancer showed negative correlation in both experiments, and DeepVelo showed negative correlation in the CHASE experiment. DeepKINET showed no negative correlation in any setting.

**Figure 3:**
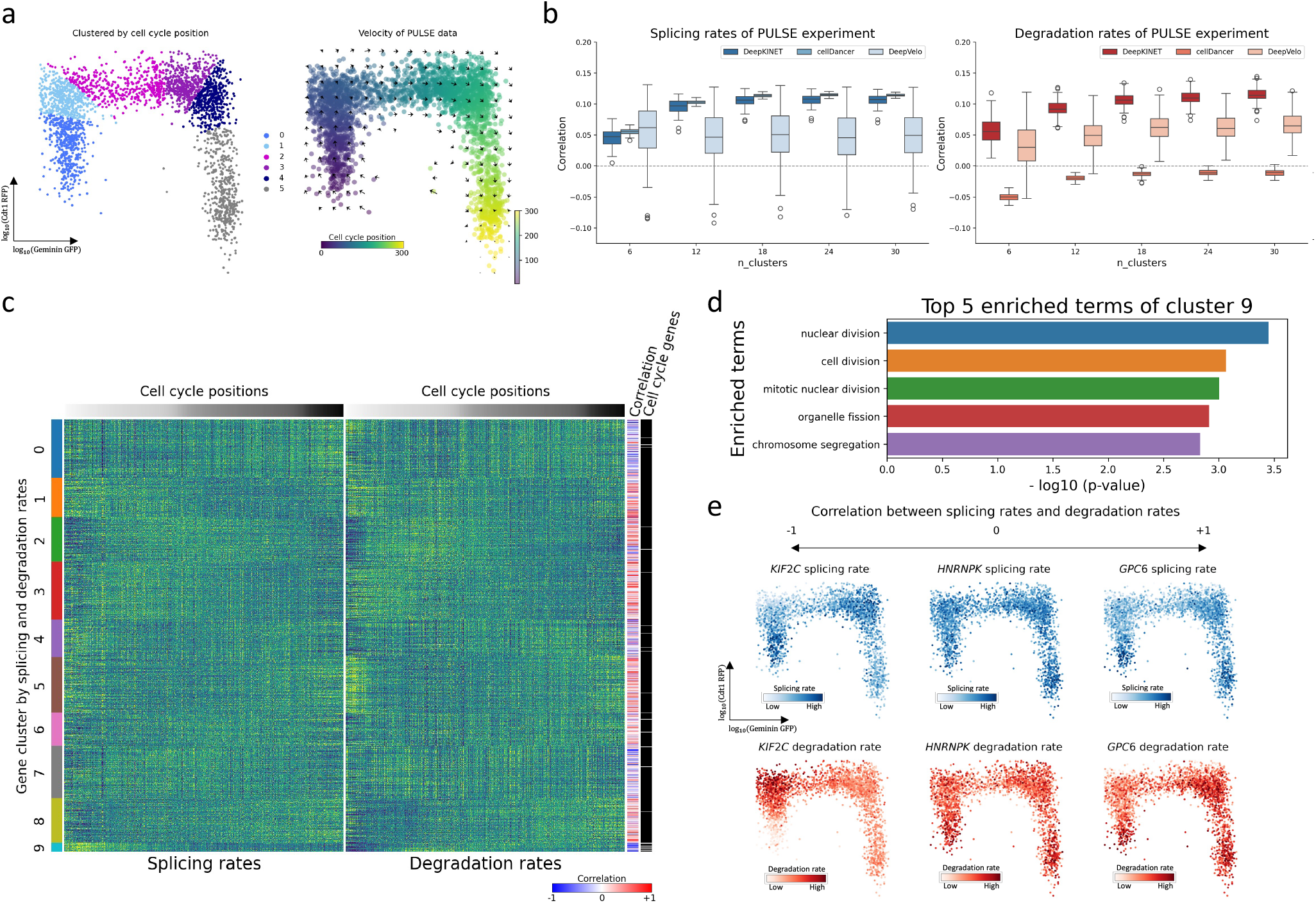
DeepKINET is also accurate for real data and outperforms cellDancer and DeepVelo. **a**. The clusters and velocities of the PULSE data were visualized on the pre-defined embedding based on the Geminin-GFP and Cdt1-RFP signals. Geminin-GFP and Cdt1-RFP signals were used in Battich et al.’s paper [Battich *et al*., 2020] to estimate cell cycle score. The cells were divided into cell clusters based on cell cycle position. The PULSE data showed an even distribution of cells with respect to cell cycle positions compared to the CHASE data (S2a). DeepKINET is able to estimate the correct direction along the cell cycle. **b**. Box plot of correlation coefficient averages between estimated rates by Dynamo and estimated rates by DeepKINET, cellDancer and DeepVelo using the PULSE experimental data. A total of 100 estimations were performed by each of DeepKINET, cellDancer, and DeepVelo. DeepKINET showed positive correlations, outperformed cellDancer and DeepVelo in terms of accuracy in degradation rates, and demonstrated comparable performance in accuracy in splicing rates. cellDancer showed negative correlations in degradation rates. **c**. Heatmaps of splicing rates (left) and degradation rates (right). To prevent extreme values from affecting the visualization, the minimum or maximum value of the top 1% was forced to the 1% and 99% quantile values. The genes were clustered by splicing and degradation rates and sorted by their clusters. The cells were sorted by cell cycle positions. The correlation coefficients between splicing and degradation rates for each gene are indicated by colored bars. The genes related to the cell cycle [Tirosh *et al*., 2016] are also shown in white color. Cluster 9 has a large number of genes related to the cell cycle. **d**. Gene Ontology (GO) terms enriched in the gene list belonging to cluster 9 obtained by g:Profiler. One-thousand genes in this analysis were used as background. **e**. Genes with different correlations between splicing and degradation rates. DeepKINET can extract genes by the value of the correlation between splicing and degradation rates. The minimum or maximum value of the top 1% was forced to the 1% and 99% quantile values.

Using the PULSE experimental data, we estimated the splicing and degradation rates for each cell and clustered the genes using these rates (Fig. 3c). We then derived the correlation coefficients between the splicing and degradation rates. Genes related to the cell cycle were concentrated in one cluster, and related terms were detected using Gene Ontology (GO) analysis (Fig. 3d). Finally, we classified the genes using the correlation coefficients between splicing and degradation rates (Fig. 3e).

Additionally, we evaluated the accuracy of DeepKINET using another metabolic labeling dataset. Single-cell metaboli-cally labeled new RNA tagging sequencing (scNT-seq) [Qiu *et al*., 2020] enables the estimation of transcription and degradation rates by distinguishing between old and new transcriptomes in the same cell. We applied DeepKINET and other methods to the hematopoiesis dataset [Qiu *et al*., 2022]. DeepKINET estimated the differentiation trajectory of hematopoietic cells, which was consistent with the results in the Dynamo paper (Fig. S3a). We applied Dynamo to estimate the degradation rates of two cell populations (Fig. S3b) and compared these estimates with those obtained from other methods. The degradation rates estimated by DeepKINET correlated with the values estimated by Dynamo and demonstrated superior performance compared to cellDancer and DeepVelo (Fig. S3c). These results further support the accuracy of DeepKINET’s estimations.

### DeepKINET to investigate functions of RNA-binding proteins and RNA-binding proteins that regulate gene clusters

We applied DeepKINET to a forebrain dataset [La Manno *et al*., 2018] to examine the functions of RNA-binding proteins. DeepKINET can classify genes based on their kinetic rates and identify RNA-binding proteins that govern these clusters. Additionally, DeepKINET can determine whether an RNA-binding protein regulates the splicing or degradation of its target genes.

First, we confirmed that the direction of RNA velocity estimated by DeepKINET was consistent with the known trajectories of cell differentiation (Fig. 4a). We then used DeepKINET to estimate the single-cell splicing and degradation rates and used these rates separately to cluster the genes. By clustering genes using either splicing rates or degradation rates independently, it becomes possible to discern which process an RNA-binding protein contributes to, thereby providing insights into its functional role in post-transcriptional regulation. We examined whether the gene clusters by kinetic rates matched the gene list of RNA-binding protein targets using Fisher’s exact test (Fig. 4b). We found clusters that matched the target gene lists, indicating that genes regulated by the same RNA-binding protein have similar splicing and degradation rate changes.

**Figure 4:**
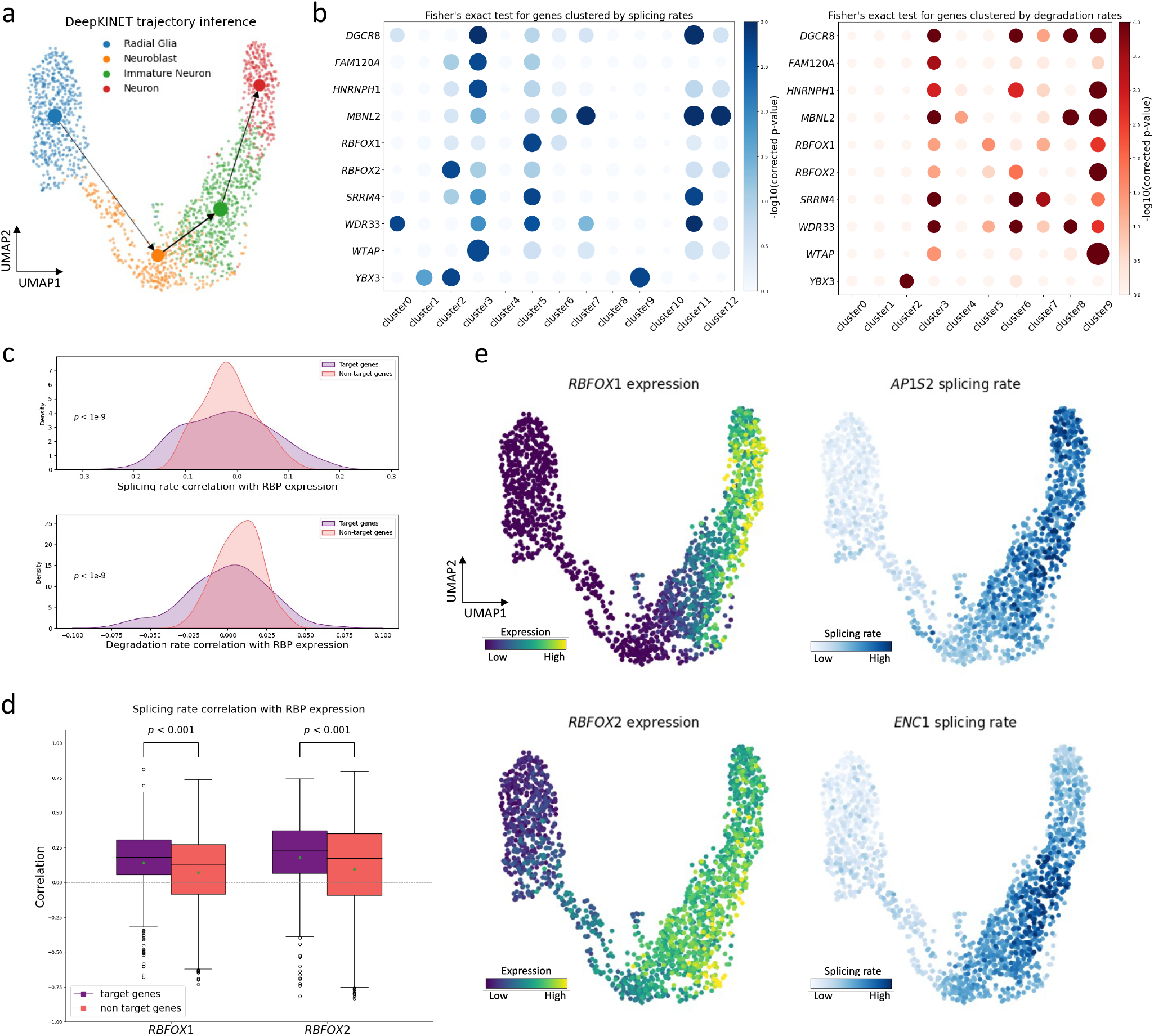
RNA-binding protein analysis of the forebrain dataset by DeepKINET. **a**. PAGA trajectory inference of forebrain dataset using DeepKINET’s velocity estimates. **b**. Dot heatmap showing the association of each RNA-binding protein (RBP) targets with each gene cluster. The genes were clustered using splicing and degradation rates separately, and Fisher’s exact test was used to determine if a list of RNA-binding protein target genes were enriched in a particular cluster. The colors indicate the corrected p-values for Fisher’s exact test. The circle size indicates the ratio of the proportion of RNA-binding protein targets in the cluster to the proportion of RNA-binding protein targets in all genes. **c**. Joint plot of the mean correlation coefficient between RNA-binding protein expression levels and the splicing and degradation rates of each target or non-target. Compared with non-target genes, target genes have higher correlations with the expression of RNA-binding proteins. A significant difference was indicated by the Levene’s test. **d**. Box plots show correlation coefficients between *RBFOX1* and *RBFOX2* expression and the splicing rates of each target or non-target gene. The green dot represents the average value. A significant difference was indicated by the one-sided unpaired t-test. **e**. Visualization of the UMAP embedding of the expression of *RBFOX1* and *RBFOX2* and the splicing rates of target genes that are highly correlated with *RBFOX1* and *RBFOX2* expression.

Next, we examined the relationship between the expression levels of each RNA-binding protein and the splicing and degradation rates of the target genes. We calculated the average correlation coefficients for both target and non-target genes for all remaining RNA-binding proteins from expression preprocessing and the observed absolute values of correlation coefficients of target genes were significantly greater than those of non-target genes (Fig. 4c). This suggests that DeepKINET accurately reflects the regulatory roles of RNA-binding proteins with respect to their target genes. Further analysis of the highly variable genes that substantially affected the kinetic rates of their targets revealed that the expression levels of *RBFOX1* and *RBFOX2* correlated with the target splicing rates (Fig. 4d), which is in agreement with established research identifying *RBFOX1* and *RBFOX2* as regulators of mRNA splicing [Conboy *et al*., 2017]. Therefore, DeepKINET demonstrated proficiency in deducing the contributions of RNA-binding proteins to splicing and degradation within the dataset, as well as in identifying genes that are potentially regulated by specific RNA-binding proteins (Fig. 4e).

### DeepKINET reveals heterogeneity in cancer cell populations

Next, we applied DeepKINET to breast cancer data to identify genes with significant changes in kinetic rates and RNA-binding proteins that exhibit distinct functions across different cell populations. Previous studies have highlighted the critical roles of splicing and degradation abnormalities in cancer development and progression [Bradley *et al*., 2023, Fang *et al*., 2022]. Additionally, the significant involvement of RNA-binding proteins in cancer has been well documented [Pereira *et al*., 2017, Qin et al., 2020]. Cell Ranger [Zheng *et al*., 2020] and Velocyto [La Manno *et al*., 2018] were used to create matrices of the spliced and unspliced breast cancer data [Liu *et al*., 2022].

We applied DeepKINET to malignant epithelial cells from the breast cancer data (Fig. 5a) and confirmed that the estimated velocities were in the direction from primary cells to metastatic cells (Fig. S4a). We then estimated the single-cell kinetic rates and identified genes that exhibited marked differences in their splicing or degradation rates when primary cells were compared with metastatic cells (Fig. 5b, c). Among these, *KDM6A* [Xiao *et al*., 2022], *PGR* [Fowler *et al*., 2020], *PIK3CA* [Fusco *et al*., 2021], *PRKAA1* [Yi *et al*., 2020], *TPM2* [Zhang *et al*., 2018], *TP63* [Gatti *et al*., 2019], *USP9X* [Guan *et al*., 2022], and *TIMP2* [Peeney *et al*., 2020] have been implicated in breast cancer metastasis. These variations in the kinetic rates may play a pivotal role in metastasis.

**Figure 5:**
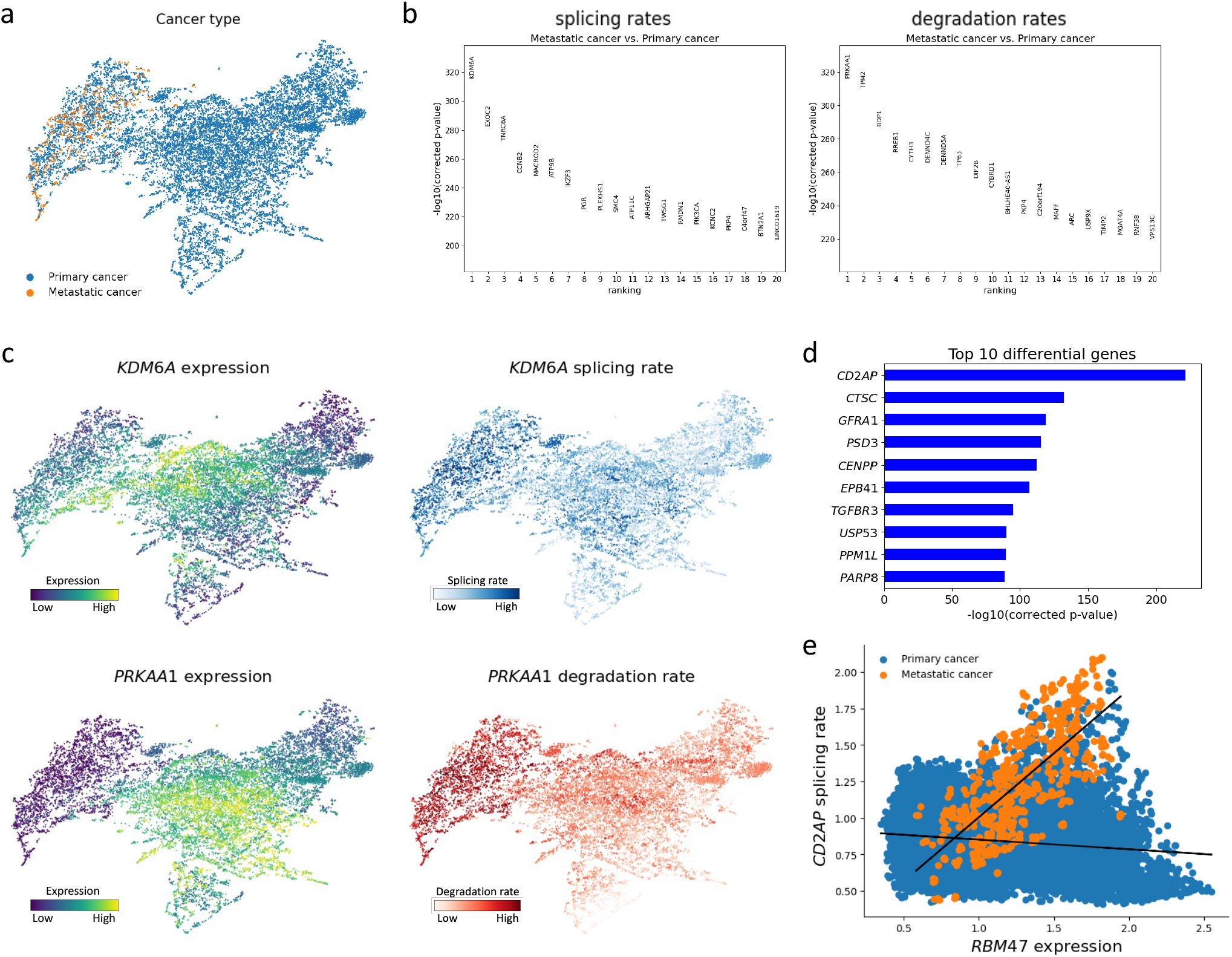
DeepKINET can identify kinetics changes involved in metastasis using breast cancer data. **a**. Visualization of UMAP embedding of the velocities estimated by DeepKINET and pre-defined classifications for malignant epithelial cells of breast cancer. There are 15,269 primary cells and 642 metastatic cells. The velocities indicate the direction from the primary cancer to the metastatic cancer. **b**. Genes with large changes in their splicing rates (left) and degradation rates (right) between primary and metastatic cells as determined using *t*-test. These genes include those involved in cancer metastasis and breast cancer. **c**. Visualization of UMAP embedding of expression levels and kinetic rates of the genes with the largest changes in their splicing or degradation rates. These genes are involved in breast cancer metastasis. To prevent the effect of extreme values in the visualization, the minimum or maximum value of the top 1% was forced to the 1% and 99% quantile values. **d**. Bar plot of corrected p-values for genes whose slopes changed significantly between primary and metastatic cells when linear regression was performed using *RBM47* expression levels and target splicing rates. Several of these genes are involved in breast cancer metastasis and metastasis of other cancers. **e**. Scatter plot of *RBM47* expression and splicing rates of *CD2AP*, the gene with the most slope change.

Furthermore, we explored the correlation coefficients between the expression of highly variable RNA-binding proteins and the kinetic rates of their target genes. Within this dataset, the effect of *RBM47* on the splicing rate of its target genes was significant (Fig. S4b, S4c). Because *RBM47* is involved in RNA splicing and metastasis, including that of breast cancer [Kim *et al*., 2019, Vanharanta et al., 2014, Guo et al., 2022], this result indicates the capacity of DeepKINET to accurately reflect authentic biological processes. We also investigated whether the relationship between *RBM47* and its target genes differed significantly between primary and metastatic cells. We performed linear regression on the expression of *RBM47* and the splicing rates of its targets, and examined whether the slope of the regression varied significantly between primary and metastatic cells. We corrected the p-values using multiple testing corrections and extracted significantly altered genes (Fig. 5d, e). Among these genes, *CTSC* [Xiao *et al*., 2021], *PSD3* [Jin *et al*., 2021], *TGFBR3* [Jovanović *et al*., 2014], and *USP53* [Liu *et al*., 2023] are involved in breast cancer metastasis. *CD2AP* [Xie *et al*., 2020], *GFRA1* [Ma *et al*., 2020], and *EPB41* [Yuan *et al*., 2021] are implicated in the metastasis of other cancers, but no findings on breast cancer metastasis have been reported. These findings imply that changes in the effect of *RBM47* expression on the splicing rates of its target genes are associated with cellular transitions critical for cancer metastasis.

### DeepKINET reveals changes in splicing due to mutations in splicing factors

Finally, we investigated changes in target splicing rates due to mutations in a splicing factor using erythroid lineage cells. The subtype of myelodysplastic syndrome (MDS), MDS-RS (MDS with ringed sideroblasts), has mutations in the splicing factor *SF3B1* and is characterized by severe anemia and the accumulation of erythrocyte progenitor cells in the bone marrow. *SF3B1* is responsible for connecting immature mRNAs to spliceosomes, and mutations in it lead to aberrant splicing, particularly the use of alternative 3’ splice sites [Ochi *et al*., 2022], resulting in reduced standard transcripts [Shiozawa *et al*., 2018].

We extended our model using the conditional VAE framework [Kingma *et al*., 2014] to integrate and analyze multiple samples (see more details in Methods). We used data from Adema et al.’s[Adema *et al*., 2022] bone marrow mononuclear cells from seven MDS-RS patients with *SF3B1* mutations and two age-matched healthy donors. We analyzed erythroid lineage cells, which are known to be damaged by MDS-RS (Fig. 6a). The results of trajectory inference were consistent with the known erythroid differentiation process (Fig. 6b).

**Figure 6:**
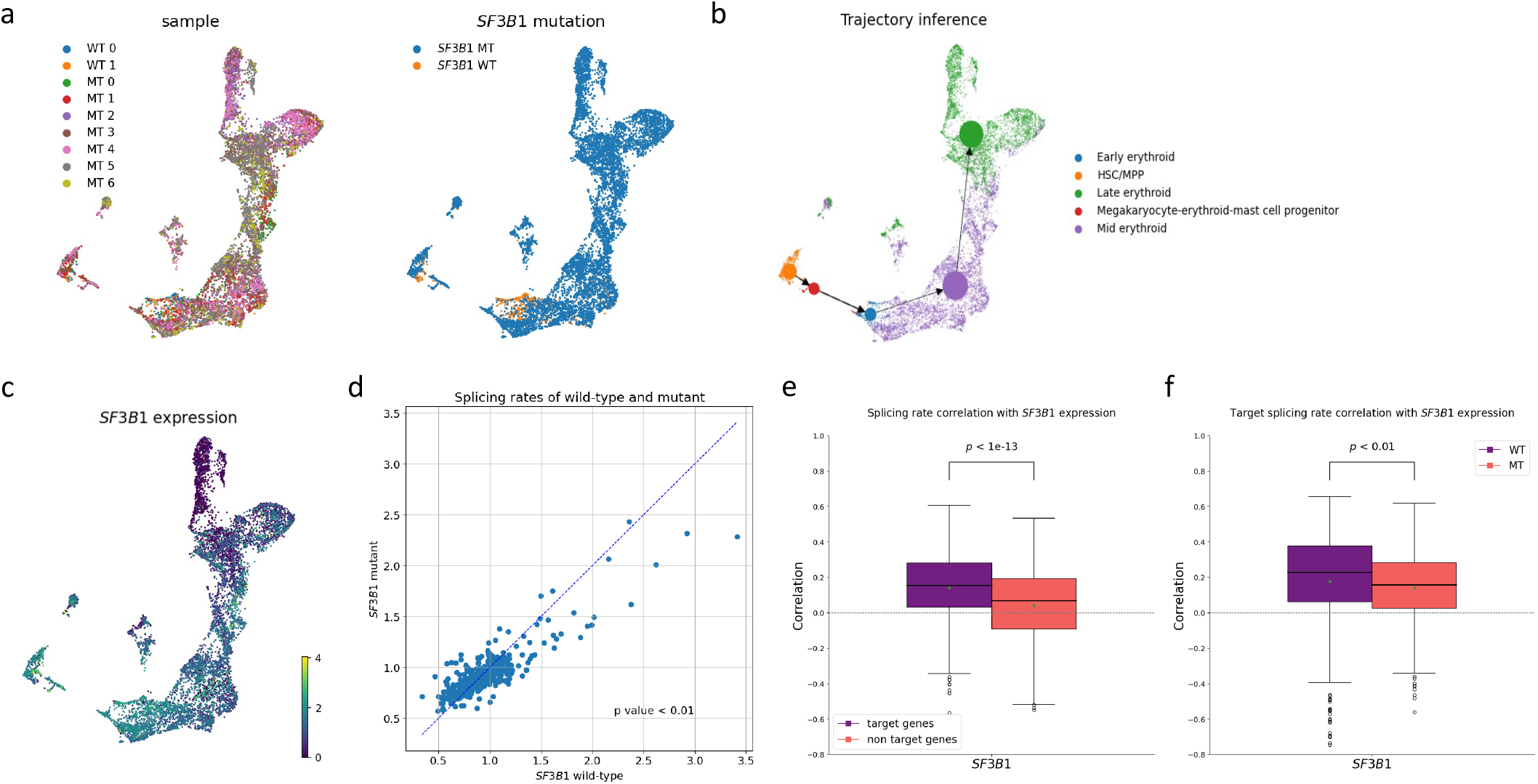
Analysis of a mutated splicing factor by DeepKINET. **a**. Patient label (left) and *SF3B1* mutation status (right) on the UMAP embedding from the low-dimensional latent cell state estimated by DeepKINET. **b**. PAGA trajectory inference using DeepKINET’s velocity estimates. DeepKINET can accurately estimate the differentiation pathway of erythroid lineage cells. **c**. UMAP coordinates colored by the expression of *SF3B1*. **d**. Scatter plot of the average splicing rate by *SF3B1* mutation status (wild-type on the x-axis and mutant on the y-axis) for each gene. A one-sided paired t-test showed that cells with the *SF3B1* mutation had significantly lower target splicing rates than cells without the mutation. **e**. Box plot showing the correlation between *SF3B1* expression and target or non-target gene splicing rates. In the target genes, the correlation value were significantly higher than those of the non-target genes. **f**. Box plot showing the correlation between *SF3B1* expression and target gene splicing rate. In cells with *SF3B1* mutations, the correlation value was significantly lower than in cells without mutations.

Our analysis revealed that the splicing rate of *SF3B1* target genes in *SF3B1* mutant cells was significantly lower than in healthy cells (Fig. 6d). This result suggests that DeepKINET effectively captures the changes in splicing kinetics caused by *SF3B1* mutations. It should be noted that in the RNA velocity model, the splicing rate is defined as the amount of change from unspliced mRNA to spliced mRNA per unit time. In the standard quantification method such as Velocyto [La Manno *et al*., 2018], reads are annotated as spliced mRNA if they map only to exon regions, and even if only a small amount maps to intron regions, they are annotated as unspliced mRNA. It is known that in the case of SF3B1 mutations, the usage of alternative 3’ splice sites results in the shifting of splicing sites tens of base pairs upstream compared to the canonical 3’ splice sites, causing an insertion of intronic sequence at the authentic exon–exon junction [Shiozawa *et al*., 2018]. From these considerations, it can be inferred that the use of alternative 3’ splicing sites in *SF3B1* mutants reduces the amount of transcripts classified as spliced mRNA, which in turn leads to decreases the splicing rates in the RNA velocity model. The results from DeepKINET suggest that this model captures the changes in kinetics underlying these biological processes.

Furthermore, the average correlation between the expression of *SF3B1* and the splicing rates of target genes was significantly higher than the average correlation between the expression of *SF3B1* and the splicing rates of non-target genes (Fig. 6e), consistent with the functional characteristics of *SF3B1*. In addition, the average correlation between the expression of *SF3B1* and the splicing rates of target genes in *SF3B1* mutant cells was significantly lower than that in *SF3B1* non-mutant cells (Fig. 6f). This results suggests that mutations in *SF3B1* make it difficult to produce the target standard spliced mRNA and that the mutated *SF3B1* does not contribute to normal splicing. These findings demonstrate that DeepKINET can capture changes in splicing of targets due to mutations in splicing factors.

## Discussion

In this study, we introduced DeepKINET, a groundbreaking method for accurately estimating splicing and degradation rates at single-cell resolution. By harnessing cell state information and RNA velocity, DeepKINET advances beyond conventional models that assign static splicing and degradation rates to genes, offering dynamic and cell-specific analysis. This innovation marks the first instance in which such kinetic rates have been estimated and validated for accuracy at the single-cell level using both simulated and metabolic labeling data, thereby enabling a more nuanced understanding of gene expression regulation. Our approach facilitates a variety of biological analyses, including clustering by the kinetic rate, identifying genes with highly variable kinetics across cell types, and detecting RNA-binding proteins that influence splicing and degradation processes. Importantly, DeepKINET utilizes readily available scRNA-seq data, avoids the need for complex metabolic labeling, and paves the way for novel investigations of gene expression kinetics. Using this method, one can gain insights into the regulatory mechanisms of gene expression and uncover potential therapeutic targets for diseases in which splicing and degradation are dysregulated, such as cancer. These insights will be critical in elucidating variations in gene expression among cells and populations, bringing to light complex regulatory networks.

Despite its advantages, DeepKINET has several inherent limitations. It employs a unified model to estimate splicing and degradation rates, which can lead to correlation trends among these rates (Fig. S1d, S2c, S5). Nonetheless, the fidelity of our estimates was supported by simulated and metabolic labeling data. In addition, the correlation between splicing and degradation rates estimated by DeepKINET was the lowest among the three methods, while cellDancer and DeepVelo exhibited high correlations. These high correlations suggest that cellDancer and DeepVelo may have limited ability to disentangle the effects of splicing and degradation. While kinetic rate estimation at the single-cell level improves the details of RNA velocity calculations [Li *et al*., 2023], the simultaneous estimation of RNA velocity and kinetic rates presents a challenge, indicating the need for further methodological enhancements and additional constraints for improved accuracy in estimating kinetic rates.

It is worth noting that by extending DeepKINET, the assumption of fixed transcription rates for each gene can also be eliminated. However, this would increase the number of parameters, and further investigation is required to determine whether the estimation of the transcription, splicing, and degradation rates would all remain stable in such a setting. An existing method MultiVelo [Li *et al*., 2023], uses multi-omics data (gene expression and chromatin accessibility) as input and assumes that transcription rates are determined based on chromatin accessibility near the gene. Considering the fact that transcription factors bind almost exclusively to open chromatin and provide dynamic regulation of transcription [Klemm *et al*., 2019], MultiVelo’s modeling is more realistic than estimating transcription rates using only scRNA-seq data and may allow for more accurate estimation of transcription rates.

A notable challenge lies in the current limitations of RNA velocity analysis in distinguishing mRNA isoforms [Gorin *et al*., 2022], with implications particularly relevant to diseases such as cancer, where alternative splicing is prevalent. Addressing this issue in future versions of DeepKINET could provide deeper biological insights and a more authentic portrayal of variations in mRNA splicing.

In summary, DeepKINET is a significant contributor to the field of single-cell biology, offering a novel analytical framework that not only advances the current understanding, but also sets the stage for future innovations that will further elucidate the complexities of cellular kinetics.

## Methods

In DeepKINET, the cell states and RNA velocity were first estimated, as in VICDYF [Nagaharu *et al*., 2022], and the learned parameters of the encoders and decoders were fixed. Subsequently, decoders are created that take the cell states as the input and output the splicing and degradation rates at the single-cell level. These decoders are trained to better reconstruct unspliced mRNA amounts.

### Derivation of single-cell splicing and degradation rates

The cell state and RNA velocity were estimated as described in the previous VICDYF method. The standard normal distribution is used as a prior for the low-dimensional latent cell state *z*_*n*_ ∈ *R*^*D*^ of cell *n* and the direction of small change *d*_*n*_ ∈ *R*^*D*^ on the low-dimensional latent cell space. *D* is the dimension of the latent cell space and the default value is 20.

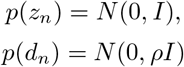

where *ρ* is a scaling factor, and *I* is the identity matrix. The direction of the small change *d*_*n*_ needs to have a small scale with respect to *z*_*n*_; thus, we set *ρ* = 0.01 to be the same as in VICDYF. Unspliced and spliced transcriptomes of a single cell are indicated by *u*_*n*_ ∈ *R*^*g*^ and *s*_*n*_ ∈ *R*^*g*^, where *g* is the number of genes. Poisson or negative binomial distributions were assumed for the distributions of *u*_*n*_ and the distribution of *s*_*n*_ given *z*_*n*_. A Poisson distribution was assumed for all analyses in this research.

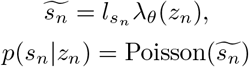

where 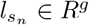 is the mean of spliced counts across all genes in the single cell, and *λ*_*θ*_(*z*_*n*_) ∈ *R*^*g*^ is the decoding neural network of the latent cell states with 100 hidden units, one hidden layer, and layer normalization. 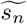is the reconstructed spliced mRNA counts. We derived the approximate time change in the mean parameter of the spliced transcriptome by decoding a small change in the latent cell state. In VICDYF, only *s* is used as input for the VAE to quantify the uncertainty of *u* given *s*. However, in DeepKINET, both *u* and *s* are used as inputs because we do not focus on the uncertainty of *u*. Moreover, to determine the small change in *s*, we differentiate the decoder transformation from *z* to 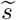 by *z* using a functorch instead of using the central difference approximation in VICDYF.

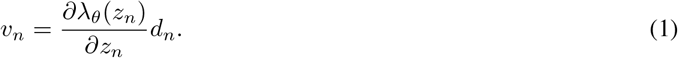

Here, we assumed that the mean parameter of the abundance of spliced and unspliced transcriptomes was represented by the differential equation of splicing kinetics as an RNA velocity estimation.

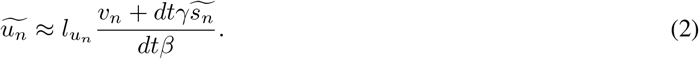

where *β* ∈ *R*^*g*^ is a vector of gene-specific splicing rates of unspliced transcripts and *γ* ∈ *R*^*g*^ is a vector of gene-specific degradation rates of spliced transcripts. Here, *β* and *γ* are the same value for each cell. 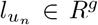 is the mean of unspliced counts across all genes in the single cell. 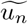is the reconstructed unspliced mRNA counts. By combining (1) and (2), we can approximate the mean parameter of the abundance of unspliced transcripts as follows:

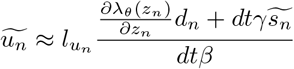

We assumed that the abundance of unspliced transcriptomes *u* has a Poisson distribution, as follows: where *dt* is the small interval and is set to 1.

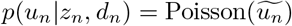

We assume the variational posterior distribution of *z*_*n*_ is a Gaussian distribution that depends on the raw counts of spliced and unspliced mRNA and the variational posterior distribution of *d*_*n*_ is a Gaussian distribution that depends on *z*_*n*_.

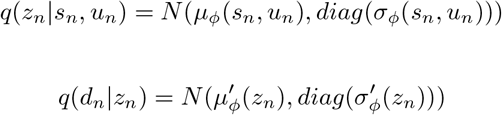

where *μ*_*ϕ*_() and *σ*_*ϕ*_() are the encoding neural networks with 100 hidden units, two hidden layers, and layer normalization [Ba *et al*., 2016]. *μ*^′^_*ϕ*_(*s*_*n*_, *u*_*n*_) and *σ*_*ϕ*_′ (*s*_*n*_, *u*_*n*_) are the encoding neural network with 100 hidden units, one hidden layers, and layer normalization.

The generative model and variational posterior distribution were optimized by minimizing the following loss function: Minimizing this loss function is equivalent to maximizing the variational lower bound (ELBO) of transcriptome distribution. This minimization allowed us to learn about the variational autoencoder of the spliced transcriptome, RNA velocity, and the reconstruction of the unspliced transcriptome.

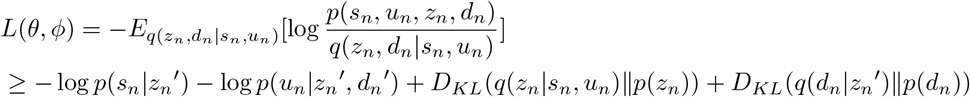

where *z*_*n*_^′^ and *d*_*n*_^′^ are derived through reparametrized sampling from *q*(*z*_*n*_| *s*_*n*_, *u*_*n*_) and *q*(*d*_*n*_ | *z*_*n*_^′^), and *E*_*p*(*x*)_[*f* (*x*)] represents the expectation of *f* (*x*) given *x*∼ *p*(*x*). The encoding neural network had 100 hidden units, two hidden layers, and layer normalization. To minimize the loss function, the Adam W optimizer was used with a learning rate of 0.001 and a mini-batch size of 100. Learning ended when the average loss of the 10 epochs was not been updated for 10 epochs.

After learning the VAE and RNA velocity, and reconstructing the unspliced transcripts as described above, all encoder and decoder parameters were fixed. Next, we create decoders that take latent variables as inputs and output splicing rate *β*_*n*_ and degradation rate *γ*_*n*_ at the single cell level. When reconstructing unspliced transcripts, they were substituted for the previous splicing and degradation rates. By estimating the splicing and degradation rates as cell-state-dependent values, the rates for cells with similar cell states will be similar, weakening the indeterminacy of the solution.

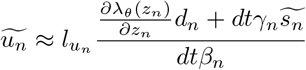

The same loss function described above was used to learn the splicing and degradation rates at the single-cell level.

### Conditional model that handles multiple samples

To address data with multiple samples, we extended DeepKINET with a conditional VAE framework [Kingma *et al*., 2014]. The prior distribution of *z*_*n*_ and *d*_*n*_ is assumed to be the same distribution as in the previous model. We assume a variational posterior distribution of cell state *z*_*n*_ with raw mRNA counts *s*_*n*_, *u*_*n*_ and batches *b*_*n*_ ∈ {0, 1} ^*B*^ as inputs. *B* is the total number of the experimental batches and *b*_*n*,*k*_ = 1 denote cell n belongs to experimental batch k.

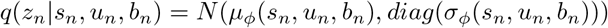

We then reconstruct spliced mRNA as follows.

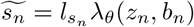

The rest of the model is the same as the model described in the previous section. After training the VAE and RNA velocity, we fix encoders and decoders and use the latent state and batch information of each cell as input to estimate the splicing rate and degradation rate of each cell.

### Creating simulated datasets

We used SERGIO (version 1.0.0) to generate the scRNA-seq count data with varying splicing and degradation rates per cluster. We used the DS6 differentiation process and the gene network from SERGIO. The SERGIO source code was rewritten to allow the splicing and degradation rates to change on a cluster-by-cluster basis. The base rate for each cell cluster was set by multiplying the SERGIO default splicing and degradation rate values by values sampled from a uniform distribution of 0.5 to 1.5. The base kinetic rates were then multiplied by values sampled from a uniform distribution of 0.75 to 1.25 for each cluster to establish different rates for each cluster. Each cluster contained 100 cells. In experiments with varying dropout rates, the `dropout indicator dynamics` function was used. Twenty dropout rate conditions were set with shape=1 and five increments from `percentile=0` to `percentile=95`. For the experiments in which the number of cells was varied, 13 conditions were set for the number of cells using a default dropout rate of `shape=1` and `percentile=65`. Ten datasets were created for each condition using different splicing and degradation rates. DeepKINET and cellDancer were used once for each dataset.

### Validation using metabolic labeling experimental dataset

Using the scEU-seq cell cycle dataset, we determined the splicing and degradation rates for each cluster using Dynamo [Qiu *et al*., 2022] and compared the estimates with those from DeepKINET and cellDancer. We split the cell cycle into PULSE and CHASE data and performed default gene filtering using Dynamo to extract 1000 genes. We divided each dataset into cell clusters based on the cell cycle position, with each cluster containing the same number of cells. We then modified the `dynamo.tl.recipe_kin_data` and `dynamo.tl.recipe_deg_data` functions to calculate the kinetic rates for each cluster. Using other parameters and following the default values of Dynamo, we derived the splicing and degradation rates for each cluster. We then applied DeepKINET, cellDancer and DeepVelo to the PULSE and CHASE data 100 times each and derived the correlation coefficients each time. In cellDancer, the seed value used for training was fixed in the source code, so the estimation was performed without setting this seed value.

For the scNT-seq hematopoietic dataset, we estimated the kinetic rates in two separate batches, each containing cells collected at different time points, as in the Dynamo tutorial. We filtered out genes exhibiting a high correlation (>0.7, 75 genes) between the moments of unspliced and spliced mRNA. We then compared the ratio of degradation rates between the two batches between the estimated values of Dynamo and the estimated values of DeepKINET, cellDancer, and DeepVelo. In DeepKINET, we used a conditional VAE framework to address batch effects by using the time batches as batch labels.

### Clustering by kinetic rates

The splicing and degradation rates of each cell were estimated using DeepKINET and Z-transformation. Principal component analysis was then performed using the rates. Leiden clustering was performed on the principal components to cluster the genes (Fig. 3c).

### Functional enrichment analysis

We performed gene clustering using kinetic rates on the cell cycle PULSE data. GO analysis was performed on each gene cluster (Fig. 3d). We used g:Profiler [Raudvere *et al*., 2019] for the analysis. When conducting GO term analysis, we used all genes used to estimate splicing and degradation rates as the background.

### Enrichment test of RNA-binding protein targets

Using the forebrain dataset, we performed gene clustering based on the kinetic rates. We examined whether the genes in each cluster were enriched for RNA-binding protein targets (Fig. 4b). We selected RNA-binding proteins that were included in the 1000 highly variable genes selected by preprocessing, for which eCLIP data were available in the CLIPdb [Yang *et al*., 2015]. Genes with at least one binding site in the eCLIP data were considered as targets. After performing Fisher’s exact test, we used the Benjamini–Hochberg method for multiple testing correction.

### Analysis of the relationship between expression of RNA-binding proteins and kinetic rates of their targets

As a preprocessing step, we used `scvelo.pp.filter_and_normalize()` with `min_shared_counts = 20` for the forebrain dataset and `min_shared_counts = 100` for the breast cancer dataset. To ensure accuracy, we estimated the kinetic rates of genes with high variability. When all the remaining RNA-binding proteins from the expression preprocessing were used in the analysis, the expression was averaged over the neighborhoods. For the forebrain dataset, we used `n_neighbors=30`. When analyzing only the RNA-binding proteins in the highly variable genes, we used the expression reconstructed from the latent variables. The top 1000 genes in the forebrain dataset and the top 2,000 genes in the breast cancer dataset were used as highly variable genes. When comparing the expression of a specific RNA-binding protein to its target or non-target kinetic rates, we used a *t*-test to determine any significant difference in the correlation coefficients between targets and non-targets.

### Preparation of breast cancer data

We downloaded the FASTQ files from the public data of Liu et al. We then created BAM files using Cell Ranger [Zheng *et al*., 2020]. Next, Velocyto [La Manno *et al*., 2018] was used to create count matrices for unspliced and spliced mRNA. We used EPCAM and KRT19 as markers of epithelial cells, following the method described by Liu et al. We used inferCNVpy to extract the cancer cells. Among the seven patients, cells from patient 5 were selected and used for further analysis because the other patients contained few metastatic cells or, conversely, too many metastatic cells or a low number of breast cancer epithelial cells. Because tumor epithelial cells are highly heterogeneous from patient to patient [Turashvili *et al*., 2017], we did not perform an integrated analysis. Cells with at least 100 expressed genes and at least 500 unique molecular identifier counts were used. As a preprocessing step, we used `scvelo.pp.filter_and_normalize()` with `min_shared_counts = 100` and `n_top_genes = 2000` to extract genes with high expression variability.

### Preparation of bone marrow mononuclear cell data

We downloaded the FASTQ files from the public data of Adema et al [Adema *et al*., 2022]. We then created BAM files using Cell Ranger [Zheng *et al*., 2020]. Next, we used Velocyto [La Manno *et al*., 2018] to create count matrices for unspliced and spliced mRNA. As in the original paper, cells with at least 100 expressed genes and at least 500 unique molecular identifier counts were used. We annotated cells using CellTypist [Domínguez Conde *et al*., 2022]. Then, we extracted erythroid lineage cells. As a preprocessing step, we used `scvelo.pp.filter_and_normalize()` with `min_shared_counts = 20` and `n_top_genes = 1000` to extract genes with high expression variability.

### Identification of targets differentially regulated by different cell populations

We performed the following linear regression using the expression levels of RNA-binding proteins and the kinetic rates of their targets. We then examined whether the slope of the regression line differed significantly among the cell populations.

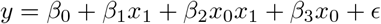

where *x*_0_ is the label of the cell population, 0 for primary cells and 1 for metastatic cells, *x*_1_ is the expression of a RNA-binding protein, and *β*_0_ to *β*_3_ are the regression coefficients. We set *β*_2_ = 0 as the null hypothesis and used `statsmodels.regression.linear_model.OLS()` to perform regression and testing. We corrected the p-values using the Benjamini–Hochberg method.

### Two-dimensional embedding of velocity

We projected the transitions in the latent space onto two-dimensional coordinates following the method described by Bergen et al [Bergen *et al*., 2020]. We used *z*_*j*_ ™ *z*_*i*_ as the change in the latent space of cell *i* to cell *j* and *d*_*i*_ as the velocity in the latent space of cell *i*. We computed a neighborhood graph, calculated the transition probabilities, and projected them onto two-dimensional coordinates using `Scvelo`’s functions.

## Acknowledgements

The computational resources for SHIROKANE were provided by the Human Genome Center at the University of Tokyo.

## Author contributions

Y.K. conceived the concept of the method. K.A. conceived the idea of validation through simulation. C.M. designed the source code for this method, conducted experiments to verify its validity, designed the analysis using this method, and performed the analysis under the supervision of Y.K. and T.S. S.N. and S.H. made minor modifications to the theory of this method. All the authors have read and approved the final version of the manuscript.

## Code and data availability

The DeepKINET implementation is available at https://github.com/3254c/DeepKINET. The modified SERGIO code used to create the simulation data is available in DeepKINET/src/SERGIO_codes. The pancreas dataset is available at `scvelo.datasets.pancreas()`, the cell cycle dataset is available at `dynamo.dataset.cellcycle()`, and the forebrain dataset is available at `scvelo.datasets.forebrain()`. Breast cancer data, including the raw data, are publicly available and can be obtained with the accession number GSE167036.

---

**Supplementary Figure 1:**
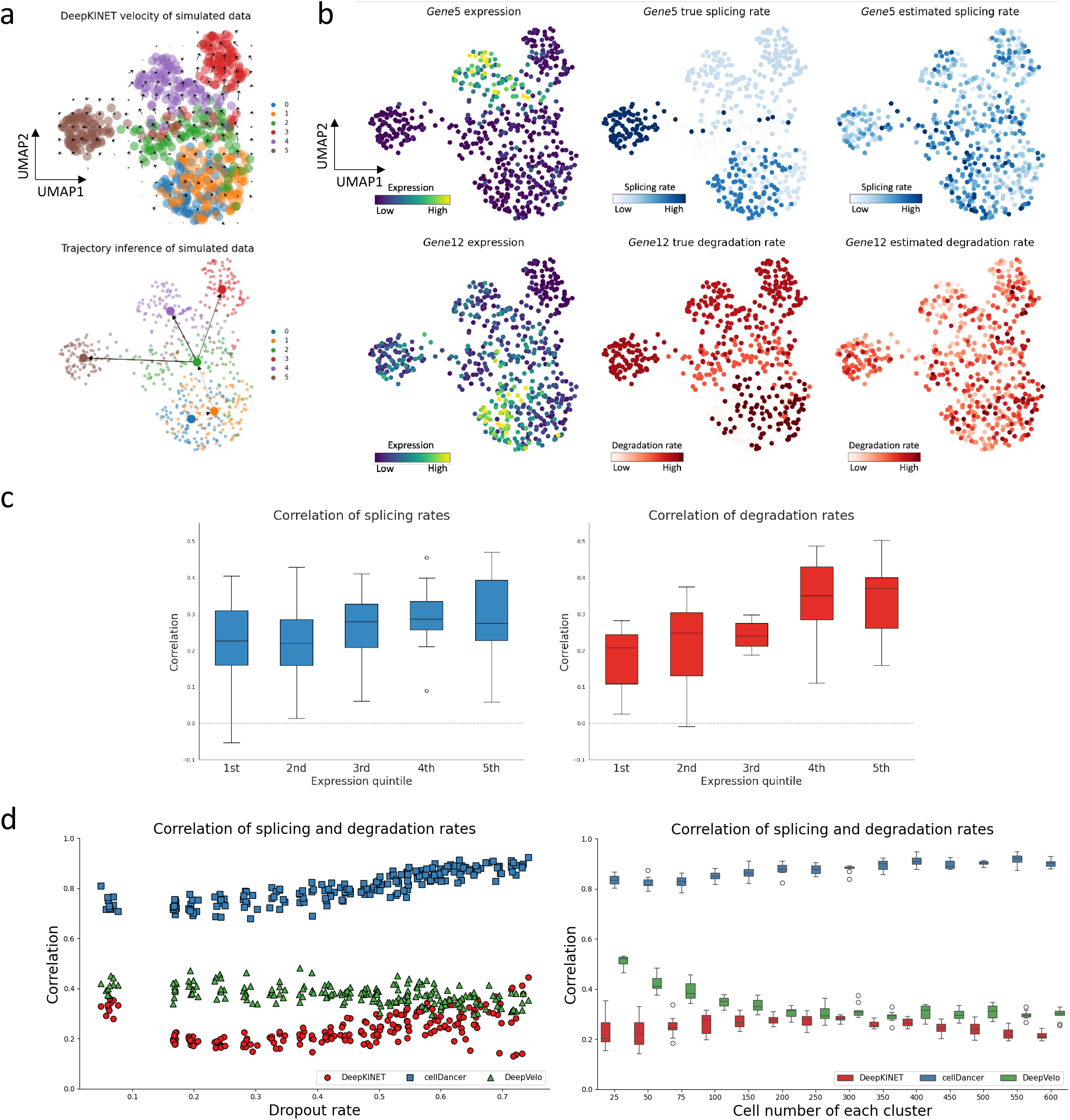
Detailed analysis of estimates from simulated data. **a**. Visualization of the UMAP embedding of the velocities (top) and PAGA trajectory inference (bottom). Each cluster contained 100 cells. **b**. Visualization of UMAP embedding of expression and splicing rates of a gene with the lowest correlation in splicing rates (top) or degradation rates (bottom). To prevent extreme values from affecting the visualization, the minimum or maximum value of the top 1% was forced to the 1% and 99% quantile values. **c**. Box plot of correlation coefficient averages when genes are separated by the sum of their expression. 1st quintile was the lowest and 5th was the highest expression. We applied DeepKINET once for each of the 10 data generated under the default dropout rate condition. **d**. The average correlation between splicing rate and degradation rate for each method. These results show that DeepKINET shows the lowest correlation in most settings and can estimate splicing and degradation rates independently. On the other hand, cellDancer has a high correlation between splicing rate and degradation rate, suggesting difficulty in optimizing both rates separately.

**Supplementary Figure 2:**
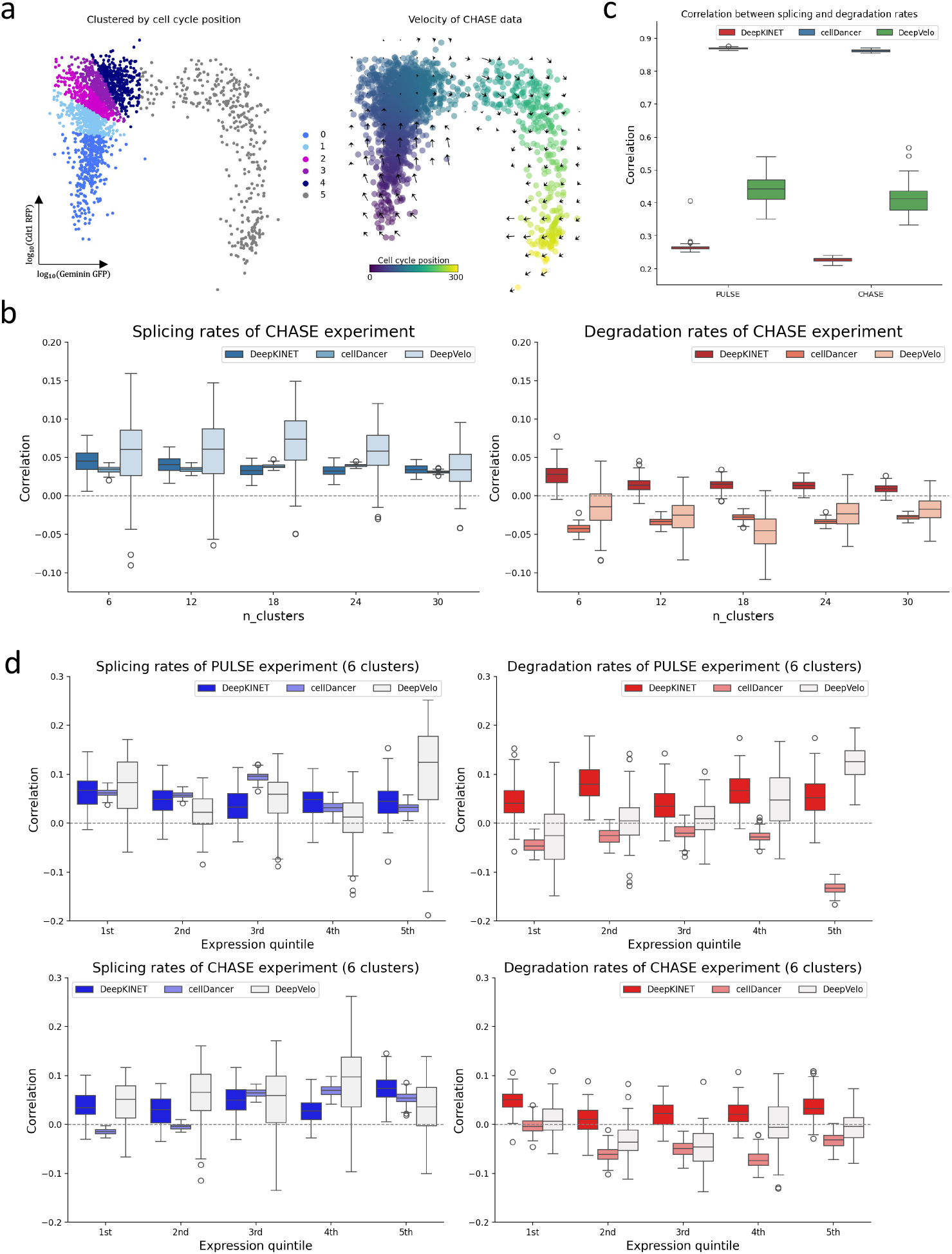
DeepKINET provides accurate kinetic rates in the CHASE data. **a**. Clusters and velocities of the CHASE dataset were visualized on the pre-defined embedding based on the Geminin-GFP and Cdt1-RFP signals. DeepKINET can estimate the correct direction throughout the cell cycle. Clustering by equivalent number of cells. CHASE data are biased for cell density. **b**. Box plot of correlation coefficients between estimated rates by Dynamo and estimated rates by DeepKINET, cellDancer and DeepVelo using the CHASE experimental data. A total of 100 estimations were performed by each of DeepKINET, cellDancer, and DeepVelo. DeepKINET showed positive correlations, outperformed cellDancer and DeepVelo in terms of accuracy in degradation rates, and demonstrated comparable performance in accuracy in splicing rates. cellDancer and DeepVelo showed negative correlations in degradation rates. **c**. The average of the absolute correlation between splicing rates and degradation rates for each method. cellDancer and DeepVelo have higher correlations than DeepKINET, suggesting that they jointly estimate splicing and degradation rates. **d**. Box plot of correlation coefficients separated by the sum of their expression in the six clusters. 1st quintile was the lowest and 5th was the highest expression. A total of 100 estimations were performed by each of DeepKINET, cellDancer, and DeepVelo.

**Supplementary Figure 3:**
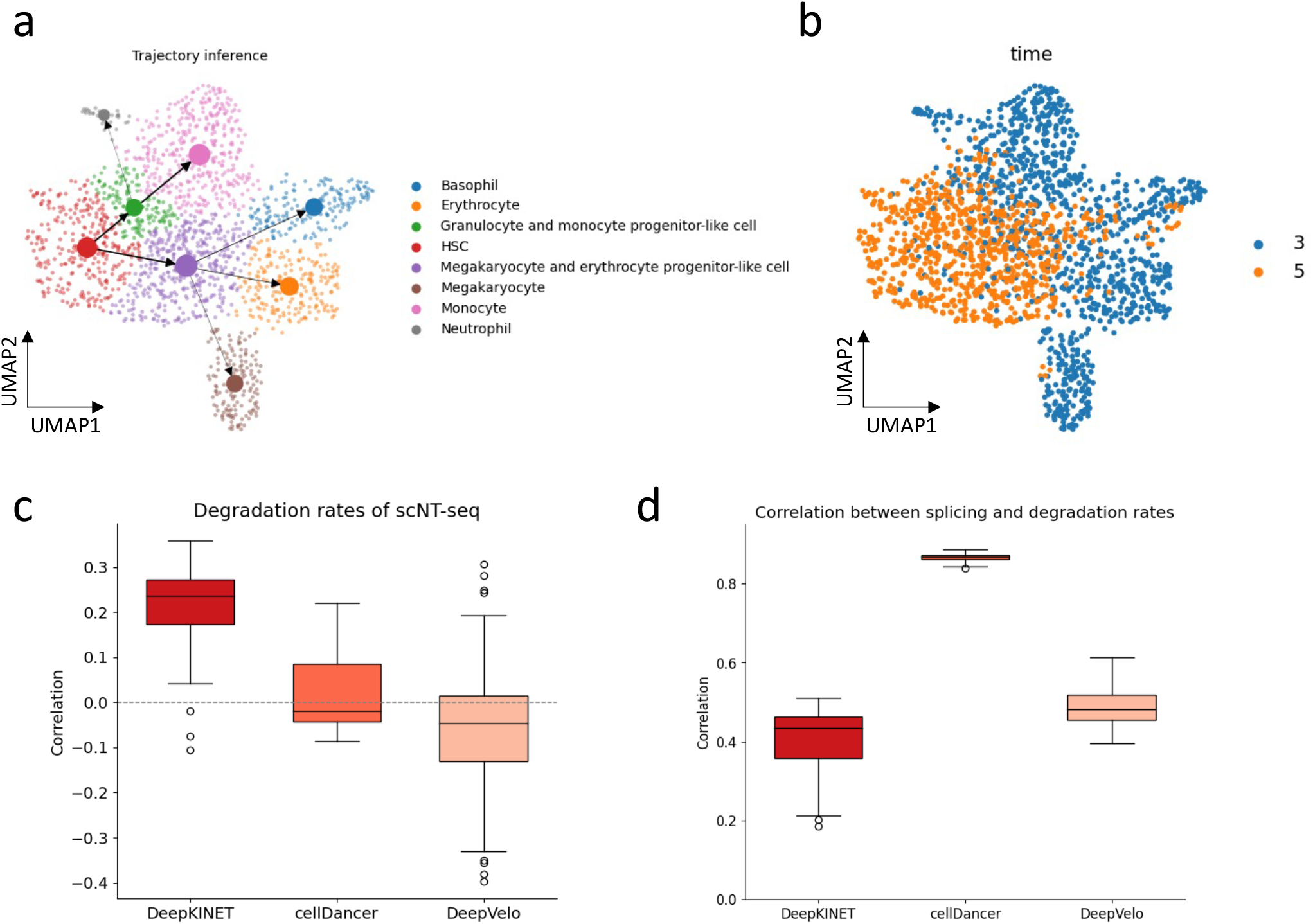
DeepKINET accurately estimates kinetic rates in the scNT-seq data. **a**. Cell type labels of the scNT-seq hematopoieisis dataset and PAGA trajectory inference using DeepKINET velocity on the predefined UMAP embedding. **b**. The time clusters of scNT-seq hematopoieisis dataset. **c**. The correlation of the ratio of degradation rates of two time clusters between Dynamo’s estimates and three estimation methods. A total of 100 estimations were performed by each of DeepKINET, cellDancer, and DeepVelo. **d**. The average of the absolute correlation between splicing rates and degradation rates for each method.

**Supplementary Figure 4:**
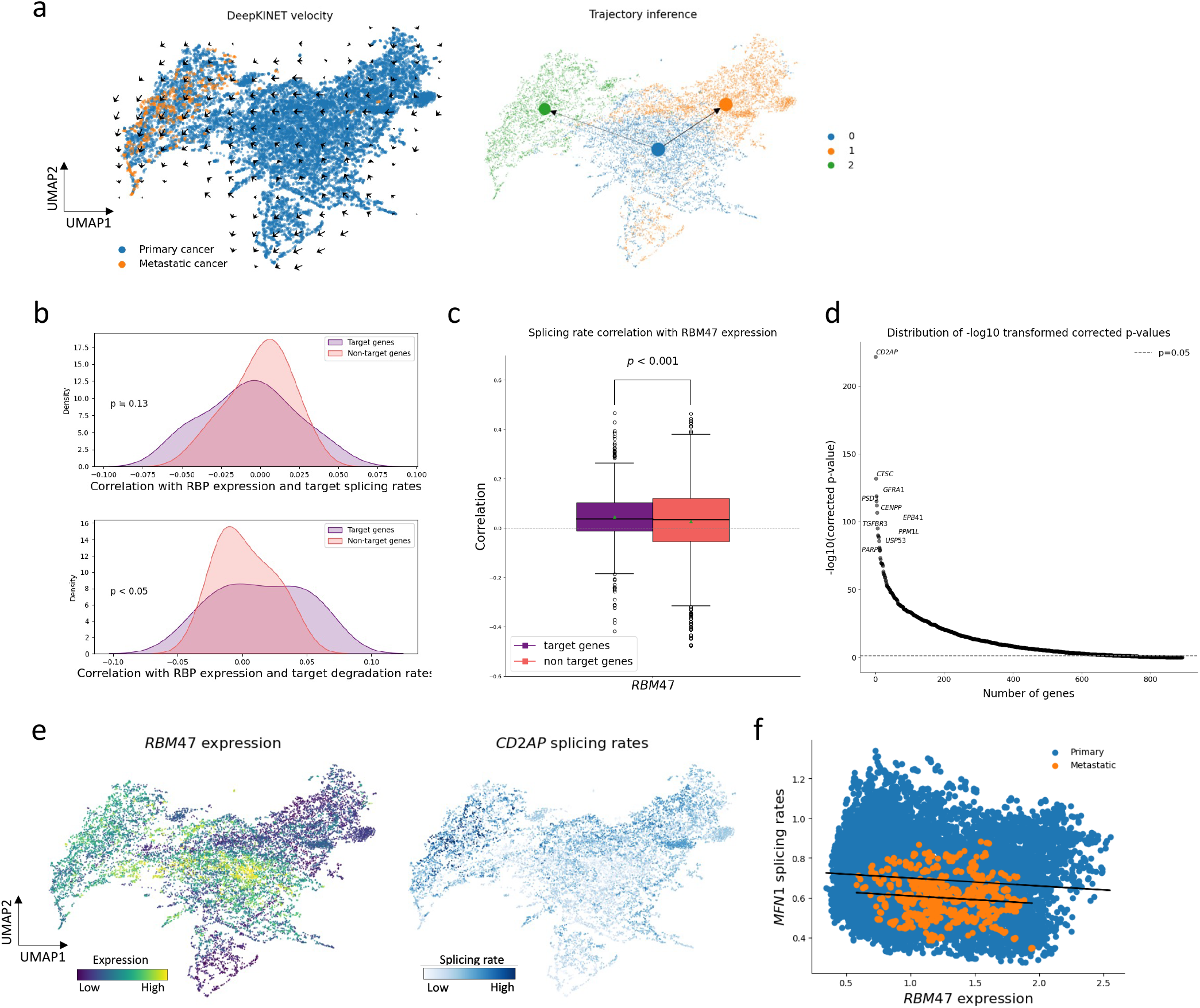
DeepKINET reveals genes differentially associated with RNA-binding proteins in primary and metastatic cells. **a**. Cancer type and velocities of the breast cancer on the UMAP embedding (left). PAGA trajectory inference using DeepKINET velocity and leiden clusters (right). The trajectory from the leiden cluster containing primary cancer cells to the leiden cluster containing metastatic cancer cells were estimated. **b**. The average correlation coefficients between highly variable RNA-binding protein (RBP) expression and the splicing and degradation rates of target or non-target. **c**. *RBM47* expression correlated with target splicing rates relative to non-target splicing rates. A significant difference was indicated by the one-sided unpaired *t*-test. The green dot represents the average value. **d**. Genes that change the slope of linear regression between primary and metastatic cells (arranged in the order of decreasing corrected p-value). **e**. Expression of *RBM47* and splicing rates of target *CD2AP* were visualized using UMAP embedding. To prevent extreme values from affecting the visualization, the minimum or maximum value of the top 1% was forced to the 1% and 99% quantile values. **f**. Scatter plot of *RBM47* expression and splicing rate of the gene with the smallest change in slope of the regression line.

**Supplementary Figure 5:**
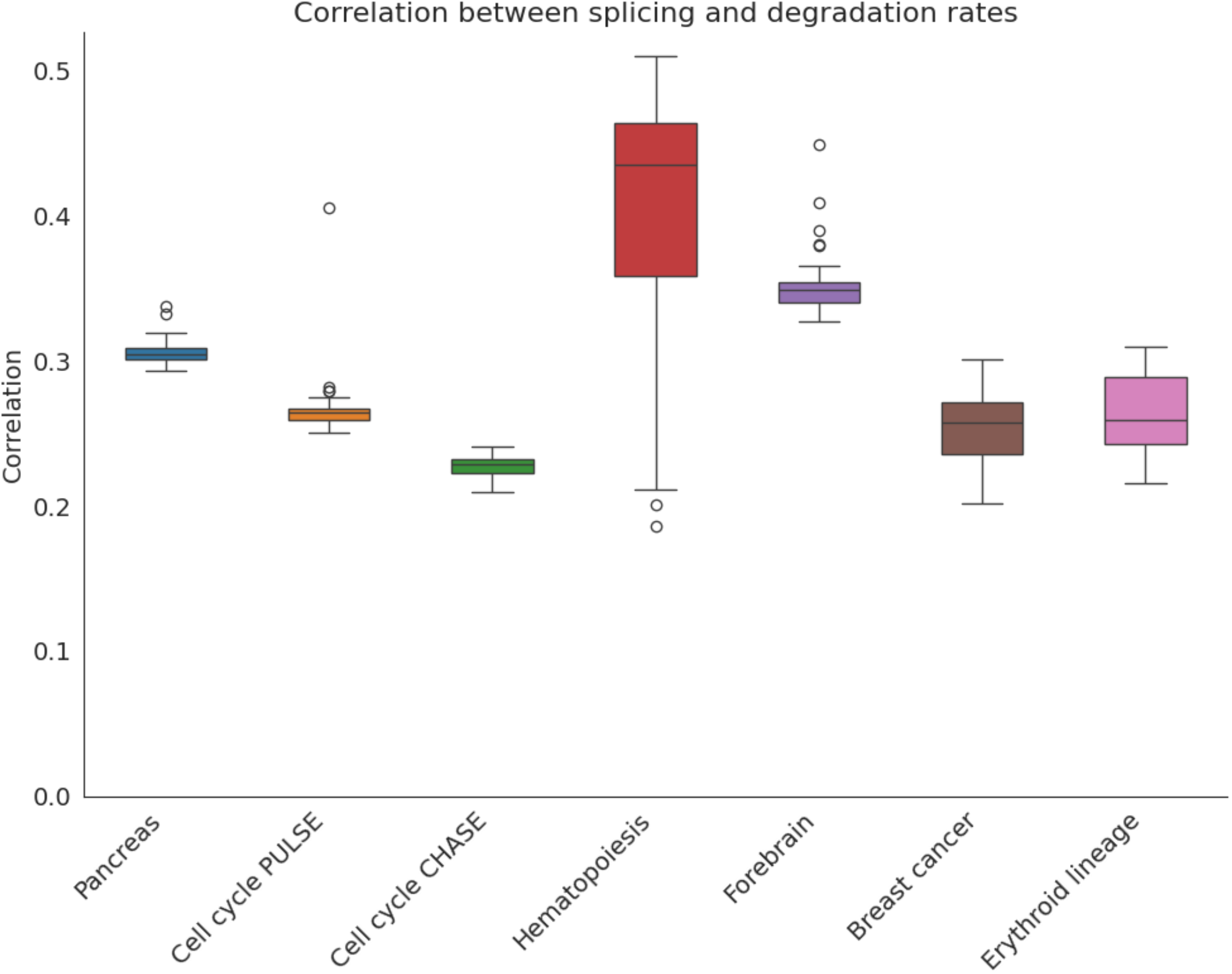
Correlation between DeepKINET estimated splicing and degradation rates. The average of the absolute correlation coefficients between splicing and degradation rates were estimated 100 times for each dataset. Splicing and degradation are weakly correlated. However, the accuracy of DeepKINET has been confirmed with simulated data and metabolic label data; thus, it captures real values. On the other hand, cellDancer and DeepKINET show higher correlations than DeepKINET (S1d, S2c, S3d).

